# Monomeric IgA antagonizes IgG-mediated enhancement of DENV infection

**DOI:** 10.1101/2021.09.14.460347

**Authors:** Adam D. Wegman, HengSheng Fang, Alan L. Rothman, Stephen J. Thomas, Timothy P. Endy, Michael K. McCracken, Jeffrey R. Currier, Heather Friberg, Gregory D. Gromowski, Adam T. Waickman

## Abstract

Dengue virus (DENV) is a prevalent human pathogen, infecting approximately 400 million individuals per year and causing symptomatic disease in approximately 100 million. A distinct feature of dengue is the increased risk for severe disease in some individuals with preexisting DENV-specific immunity. One proposed mechanism for this phenomenon is antibody-dependent enhancement (ADE), in which poorly-neutralizing IgG antibodies from a prior infection opsonize DENV to increase infection of F_c_ gamma receptor-bearing cells. While IgM and IgG are the most commonly studied DENV-reactive antibody isotypes, our group and others have described the induction of DENV-specific serum IgA responses during dengue. We hypothesized that monomeric IgA would be able to neutralize DENV without the possibility of ADE. To test this, we synthesized IgG and IgA versions of two different DENV-reactive monoclonal antibodies. We demonstrate that isotype-switching does not affect the antigen binding and neutralization properties of the two mAbs. We show that DENV-reactive IgG, but not IgA, mediates ADE in an F_c_ gamma receptor-positive K562 cells. Furthermore, we show that IgA potently antagonizes the ADE activity of IgG. These results suggest that levels of serum DENV-reactive IgA induced by DENV infection might regulate the overall ADE activity of DENV-immune plasma *in vivo* and warrants further study as a predictor of disease risk and/or therapeutic.

## Introduction

Dengue virus (DENV) is one of the most widespread vector-borne viral pathogens in the world. Consisting of four antigenically and genetically distinct serotypes (DENV-1, -2, -3, and -4), DENV is transmitted primarily by the tropical and subtropical mosquitoes *Aedes aegypti* and *A. albopictus* [1, 2]. DENV and its mosquito vectors can currently be found across Central and South America, South and South-East Asia, the Western Pacific, and sub-Saharan Africa, meaning 40% of the world’s population is currently at risk of exposure and infection [1-3]. Consequently, an estimated 400 million DENV infections are thought to occur every year, resulting in 100 million clinically apparent infection [2]. Approximately 500,000 cases per year progress to severe dengue–characterized by thrombocytopenia, vascular leakage and hemorrhage–resulting in nearly 20,000 deaths [4-7].

A distinct epidemiological feature of dengue as compared to other flaviviral diseases is the increased risk for severe disease upon heterologous secondary infection [8]. While the risk factors associated with developing severe dengue upon secondary DENV exposure are complex and incompletely understood, the leading mechanistic explanation for this phenomenon is a process known as antibody-dependent enhancement (ADE) [9, 10]. ADE is thought to occur when poorly-neutralizing or sub-neutralizing concentrations of DENV-reactive IgG opsonizes DENV and facilitates its entry into permissive F_c_γR-bearing cells [11]. Various lines of evidence support the association of ADE with severe dengue, including increased incidence of severe dengue in infants born to dengue-immune mothers [12-14]; increased viremia in interferon receptor-deficient mice or non-primates passively immunized with anti-DENV antibodies [15, 16]; and increased incidence of severe dengue during the second of sequential/heterologous DENV outbreaks and in patients with a narrow range of preexisting anti-DENV antibody titers [17, 18]. Furthermore, *in-vitro* assessments of serum ADE activity in dengue-primed non-human primates have been shown to correlate with viral titers following heterologous attenuated DENV infection [19].

The increased risk of severe dengue upon secondary heterologous infection also presents a challenge to vaccine development as incomplete or waning vaccine-elicited immunity may place recipients at an increased risk of developing severe dengue should they be exposed following vaccination [20]. This is most significantly highlighted by the revelation that the only currently US FDA licensed DENV vaccine (Dengvaxia®) fails to protect previously DENV naïve individuals from infection, and can increase the risk of hospitalization with virologically confirmed dengue [21-23]. Accordingly, understanding the subtleties of both natural and vaccine-elicited DENV humoral immunity is critical for further our understanding of disease risk and infection-associated immunopathogenesis.

To date, the literature on dengue serology has overwhelmingly focused on the contribution of immunoglobulin isotypes IgM and IgG to functional dengue immunity and infection-associated immunopathogenesis. During both primary and secondary DENV infection, these isotype antibodies follow a highly predictable pattern of induction, with an IgM response preceding the rise of DENV-reactive IgG, and DENV-reactive IgG reaching significantly higher titers during secondary infection [24, 25]. These characteristics, as well as the assumed importance of IgG-mediated ADE, have left the role of other serum antibody isotypes relatively unexamined. Notably, this includes IgA, the second most prevalent antibody isotype in serum and one that has been suggested to play a unique and non-redundant role in many viral infections [26]. Most work on DENV-reactive serum IgA has focused on its potential as a diagnostic tool [27], with a small body of literature examining DENV-reactive serum IgA as a possible correlate of severe disease [28-31].

Our group and others recently reported that IgA was the dominant isotype-switched antibody expressed by circulating plasmablasts during acute primary DENV infection [32, 33]. IgA-expressing plasmablasts were also observed in secondary dengue, but constituted a smaller fraction of the total infection-elicited immune response [32, 33]. Importantly, the IgA antibodies expressed by these plasmablasts exhibited comparable DENV-binding and DENV-neutralization activity to IgG antibodies derived from contemporaneous samples [32]. Given the milder symptoms and lower viral burden typically associated with primary dengue relative to secondary dengue, we hypothesized that DENV-reactive IgA may play some role in limiting DENV propagation and potentially the immune-mediated enhancement of diseases.

To test this hypothesis, we isotype-switched pairs of monoclonal antibodies to show that conversion of IgG to IgA does not impact the ability of a monoclonal antibody to bind whole DENV virions or to neutralize DC-SIGN-dependent DENV infection of a susceptible cell line. However, while DENV-reactive IgG antibodies exhibited potent infection-enhancing activity in *in vitro* ADE assays, we observed that DENV-reactive IgA is incapable of mediating ADE. Additionally, we observed that adding DENV-reactive monoclonal IgA to either an enhancing concentration of monoclonal IgG or to an enhancing dilution of dengue-immune plasma antagonizes ADE in a dose-dependent fashion. These results shed new light on the role of the IgA component of the humoral response to DENV, and suggest a new avenue of prophylactic and therapeutic approaches to disease.

## Results

### DENV binding and neutralizing is unaffected by antibody F_c_ isotype

To assess the potential contribution of DENV-reactive IgA to a functional anti-DENV humoral immune response, we synthesized two pairs of previously described DENV-reactive monoclonal antibodies with either an IgG or an IgA Fc domain (**Figure 1A)**. Both mAbs selected for this analysis were previously determined to bind the fusion loop of the DENV E protein and to react with all 4 DENV serotypes [32]. However, VDB33 was initially identified as an IgG clone, while VDB50 was discovered as an IgA clone (**Table 1**). This cross-conversion strategy was chosen so as to determine if the native F_c_ configuration of a given antibody influenced its functionality as either an IgG or IgA protein product.

**Table 1.**
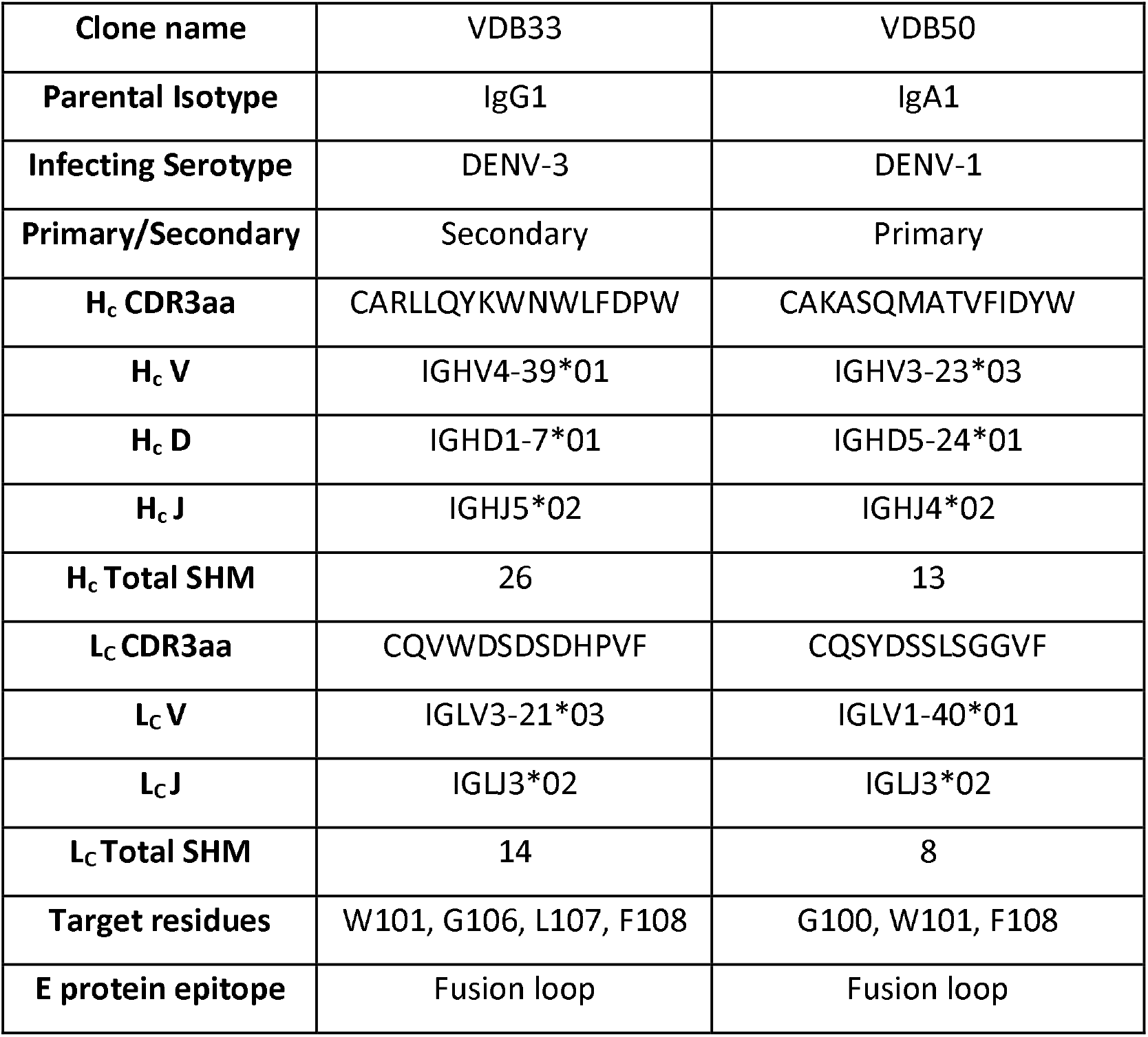
Sequence information of DENV-reactive monoclonal antibodies.

**Figure 1:**
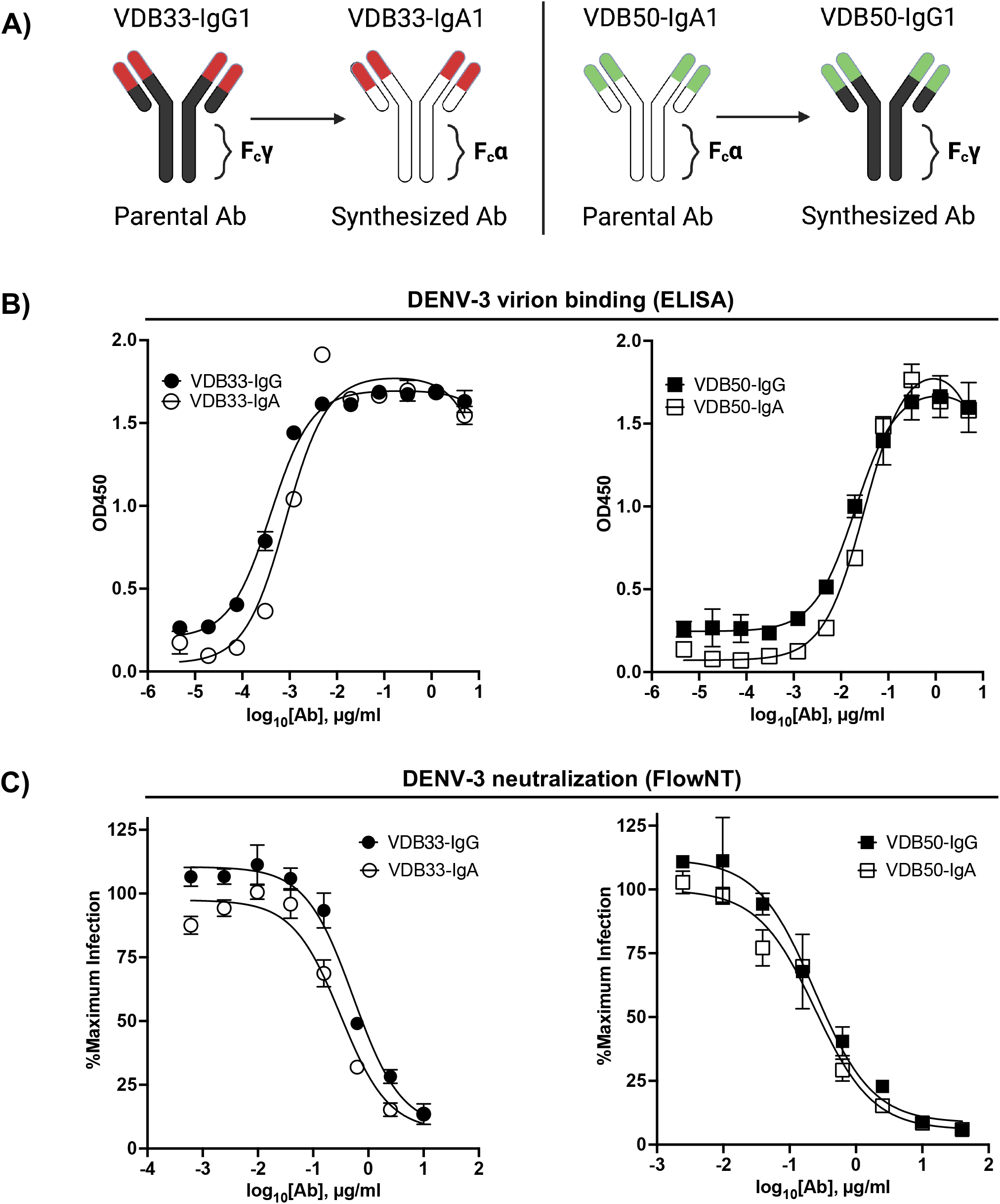
Isotype conversion scheme, DENV binding, and DENV neutralization capacity of VDB33 and VDB50 mAbs. **A)** Schematic of isotype conversion of VDB33 and VDB50 from respective parental isotypes, indicating conservation of antigen-binding domains and alteration of Fc domains. **B)** DENV-3 binding capability of VDB33-IgG, VDB33-IgA, VDB50-IgG, and VDB50-IgA measured by DENV virus-capture ELISA. **C)** DENV-3 neutralization capability of VDB33-IgG, VDB33-IgA, VDB50-IgG, and VDB50-IgA as assessed by FlowNT. Neutralization data are presented as a percent of the positive (no neutralizing mAb) control for each replicate. Error bars +/- SEM.

The DENV-binding capacity of the IgG and IgA versions of VDB33 and VDB50 was initially assessed with a DENV virion-capture ELISA. For this analysis, DENV-3 was chosen as the prototypic DENV serotype as previous work demonstrated that the IgG versions of both VDB33 and VDB50 exhibited significant DENV-3 reactivity [32]. Consistent with previously published reports, both VDB33 and VDB50 exhibited potent DENV-3 binding activity with VDB33 demonstrating ∼200 fold higher affinity for DENV-3 than VDB50 (**Figure 1B, Table 2**). However, the DENV-binding capacity of the two mAbs was not impacted by their conversion to either an IgG or IgA format (**Figure 1B, Table 2**). Furthermore, this cross-conversion of VDB33 and VDB50 to either an IgG or IgA format minimally impacted the DENV-3 neutralization activity of the clones when assessed using a flow cytometry-based neutralization assay (**Figure 1C, Table 2**). These results indicate that both IgG and IgA isotype antibodies are equally capable of binding and neutralizing DENV, reaffirming that antibody epitope/paratope interactions occur independently of an antibody’s F_c_ domain.

**Table 2.** Functional characteristics isotype-switched monoclonal antibodies.

|  |  |  |  |  |
| --- | --- | --- | --- | --- |
| **Clone name** | VDB33 IgG | VDB33 IgA | VDB50 IgG | VDB50 IgA |
| **Kd (ng/mL)** | 0.3962 | 0.8296 | 19.92 | 31.12 |
| **IC50 (ug/mL)** | 0.5339 | 0.3139 | 0.2391 | 0.2439 |

### DENV-reactive IgA is incapable of mediating ADE

Having demonstrated that the antigen binding and neutralization capacity of DENV-reactive monoclonal antibodies is negligibly impacted by the isotype of the construct, we endeavored to determine if the infection-enhancing capability of these antibodies was impacted by their isotype conversion. To this end, we utilized a K562-based ADE assay, wherein antibody/DENV immune complexes were pre-formed and added to the F_c_-receptor expressing K652 cell line to assess the ability of defined antibody complexes to enhance DENV infection.

The IgG versions of both VDB33 and VDB50 exhibited potent infection-enhancing activity in the K562 ADE assay, with both antibodies capable of facilitating DENV infection/enhancement in a dose-dependent fashion (**Figure 2A-2D**). Consistent with their relative EC_50_/IC_50_ values, VDB33-IgG exhibited notably higher ADE activity than VDB50-IgG, but with the peak of ADE activity occurring at a similar antibody concentration. However, no infection enhancement was observed when the same assay was performed with either VDB33-IgA or VDB50-IgA (**Figure 2A-2D**). This was despite the fact that these IgA isotype antibodies exhibit nearly identical virus binding and neutralization activity as their IgG counterparts, underlining the obligate role of an antibody’s F_c_ domain in determining the ADE potential of an antibody.

**Figure 2:**
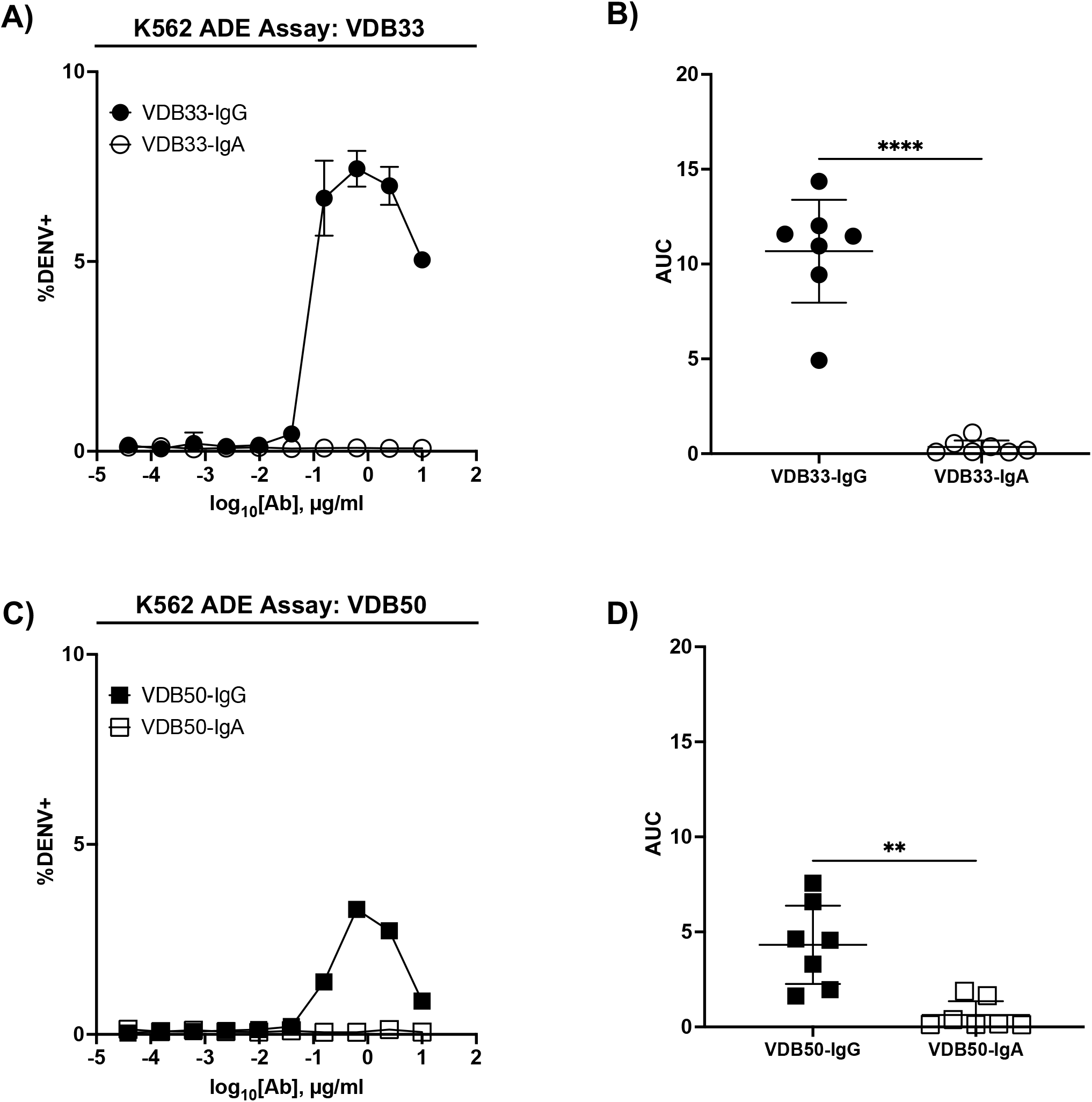
ADE activity of DENV-reactive IgG and IgA isotype antibodies. **A)** ADE activity of VDB33-IgG and VDB33-IgA against DENV-3 in K562 cells. **B)** AUC values of 7 independent experimental replicates of DENV-3 ADE assay with VDB33-IgG and VDB33-IgA **C)** ADE activity of VDB50-IgG and VDB50-IgA against DENV-3 in K562 cells. **D)** AUC values of 7 independent replicates of DENV-3 ADE assay with VDB50-IgG and VDB50-IgA. Error bars +/- SEM. ** p < 0.01, **** p < 0.0001, unpaired t test.

### DENV-reactive IgA antagonizes IgG-mediated enhancement of DENV infection

In light of the inability of VDB33-IgA and VDB50-IgA to facilitate ADE of DENV-3, we next endeavored to determine how DENV-reactive IgG and IgA behave in a polyclonal/competitive setting. IgG and IgA antibodies are never found in isolation in a dengue immune individual, so determining how these antibodies function in a complex/poly-immune setting is critical for understanding their potential contribution to function anti-DENV immunity.

To this end, we utilized the same K562 ADE assay as previously described, but used a fractional IgG/IgA replacement strategy wherein the total amount of antibody remained the same across the different titration schemes but the ratio of IgG to IgA was varied from 100:0 to 0:100. The fractional addition of DENV-reactive IgA significantly reduced the ADE activity observed in cultures containing either VDB33-IgG or VDB50-IgG (**Figure 3**). While both VDB33-IgA and VDB50-IgA were capable of antagonizing IgG-mediated ADE of DENV-3, the highly avid yet non-enhancing VDB33-IgA antibody was capable of dramatically blunting IgG-mediated ADE even when used at low fractional concentrations. Of note, the addition of DENV-reactive IgA to these ADE assays does not appear to shift the antibody dilution at which maximal ADE activity is observed for any of the cultures. Rather, the addition of DENV-reactive IgA reduces the magnitude of infection achieved at any given antibody dilution. These results are consistent with IgA actively antagonizing IgG mediated ADE by competing with DENV-reactive IgG for the same viral epitopes.

**Figure 3:**
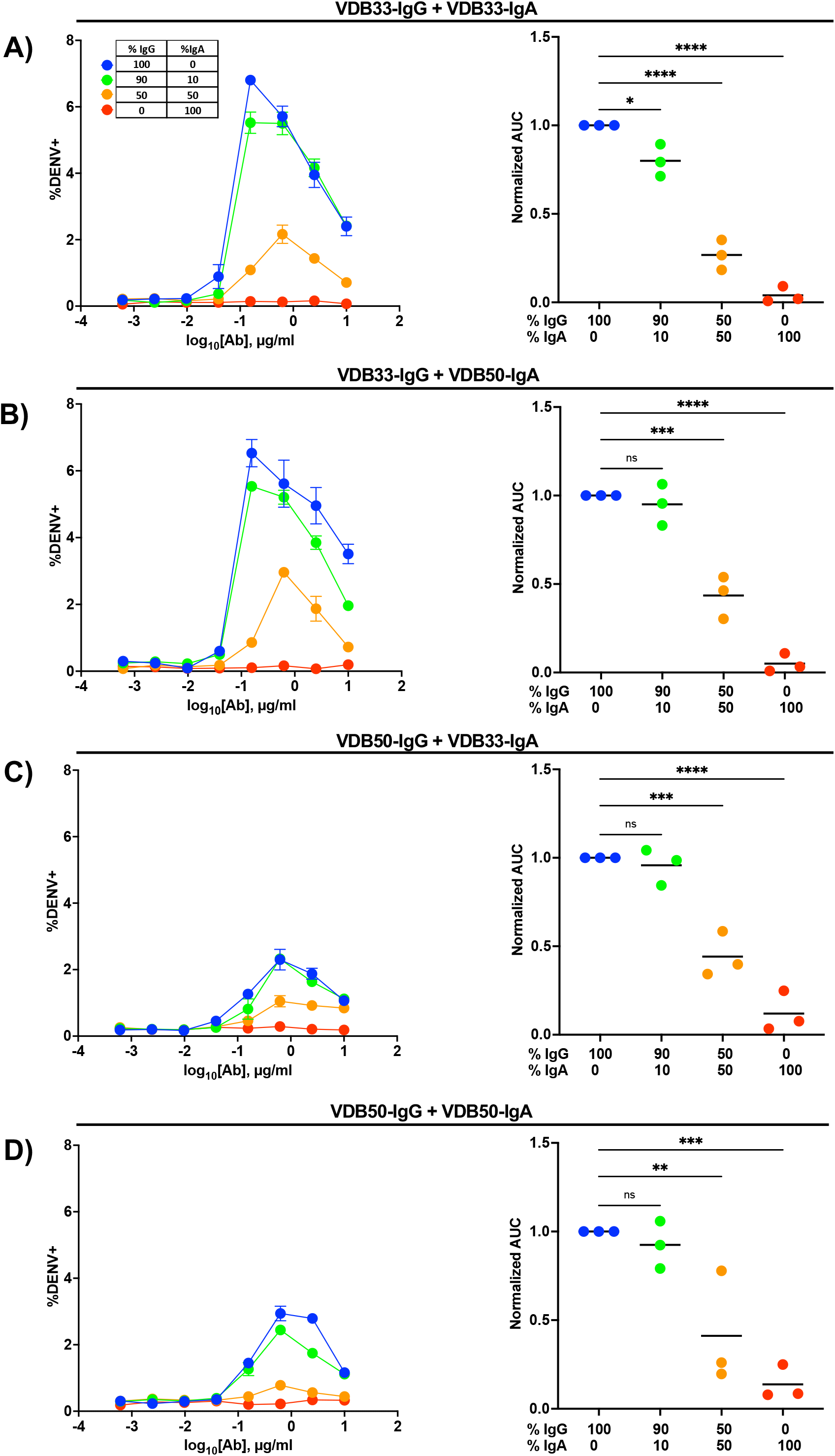
Homotypic and heterotypic monoclonal IgA antagonizes IgG-mediated antibody-dependent enhancement. **A)** DENV-3 ADE activity of VDB33-IgG when antagonized with VDB33-IgA. Total antibody concentration for each dilution point was held constant, with varying ratios of VDB33-IgG and VDB33-IgA as indicated. AUC of each ADE titration was calculated and normalized to that of the 100% IgG condition. **B)** DENV-3 ADE activity of VDB33-IgG when antagonized with VDB50-IgA. The AUC of each ADE titration was calculated and normalized to that of the 100% VDB33-IgG condition. **C)** DENV-3 ADE activity of VDB50-IgG when antagonized with VDB33-IgA. AUC of each ADE titration was calculated and normalized to that of the 100% VDB50-IgG condition. **D)** DENV-3 ADE activity of VDB50-IgG when antagonized with VDB5o-IgA. AUC of each ADE titration was calculated and normalized to that of the 100% VDB33-IgG condition. Blue = 100% IgG / 0% IgA. Green = 90% IgG / 10% IgA. Orange = 50% IgG / 50% IgA. Red = 0% IgG / 100% IgA. * p < 0.05, ** p < 0.01, *** p < 0.001, **** p < 0.0001 1-way ANOVA with Dunnett correction for multiple comparisons

### DENV-reactive IgA antagonizes DENV-immune serum mediated enhancement of DENV infection

A limitation of the analysis presented thus far is that all the monoclonal antibodies used in this analysis have the same antigen specificity; namely the fusion loop of the DENV E protein. Therefore, it is unclear what impact–if any–DENV-reactive IgA would have in the presence of a polyclonal IgG repertoire of divergent DENV antigen specificity. Therefore, we endeavored to determine how the presence of either VDB33-IgA or VDB50-IgA impacts the infection-enhancing potential of polyclonal/DENV-immune serum.

Plasma from DENV-immune donors were screened to identify samples with both high DENV-3 reactive IgG titers by ELISA as well as DENV-3 enhancing activity in the K562 ADE assay. Samples from four subjects were selected for additional analysis based on these criteria (**Figure 4A, Supplemental Figure 3, Supplemental Figure 4**). VDB33-IgA or VDB50-IgA were then titrated into cultures containing this enhancing DENV-immune plasma to determine if IgA isotype monoclonal antibodies could antagonize polyclonal enhancement of DENV-3 infection.

**Figure 4:**
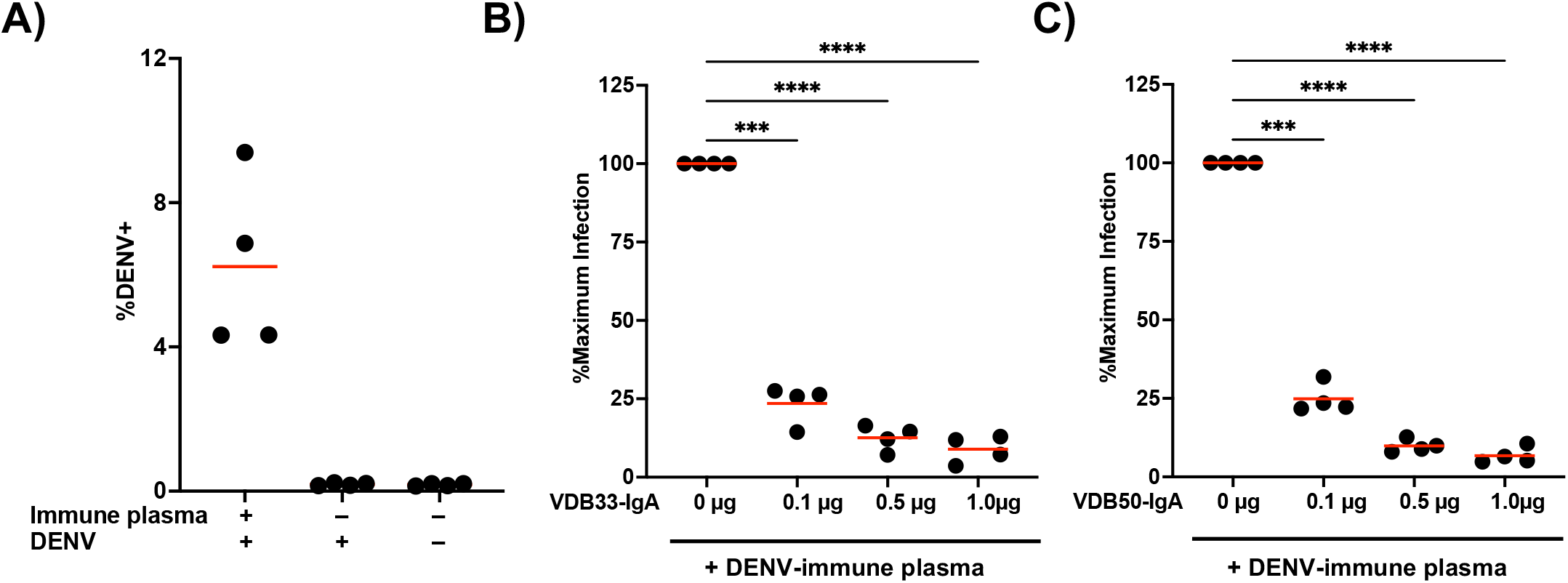
Monoclonal IgA antagonizes ADE mediated by polyclonal DENV-immune plasma. **A)** DENV immune plasma enhances DENV-3 infection of K562 cells. Each datapoint represents a unique plasma donor (n = 4). **B)** VDB33-IgA antagonizes *in vitro* enhancement of DENV-3 infection mediated by polyclonal DENV-immune serum. Serum used at a 1:50 dilution for ADE assay, n = 4 unique plasma donors. The percentage of DENV-positive cells was normalized to that observed in the plasma-only condition. **C)** VDB50-IgA antagonizes *in vitro* enhancement of DENV-3 infection mediated by polyclonal DENV-immune serum. Serum used at a 1:50 dilution for ADE assay, n = 4 unique plasma donors. The percentage of DENV-positive cells was normalized to that observed in the plasma-only condition. *** p < 0.001, **** p < 0.0001 1-way ANOVA with Dunnett correction for multiple comparisons

Consistent with what was observed with IgG monoclonal antibodies, the addition of VDB33-IgA or VDB50-IgA significantly suppressed ADE-mediated K562 infection with DENV-3 (**Figure 4B, Figure 4C**). The additional of DENV-reactive IgA in these assays suppressed ADE-mediated infection by 75%-90% in a dose-dependent fashion, a result consistent with the concept that IgG antibodies targeting the fusion loop of the DENV E protein are particularly amenable to facilitating ADE activity and are abundant in DENV-immune serum [34, 35]. These data also indicate that even modest concentrations of DENV-reactive IgA can significantly antagonize polyclonal IgG-mediated enhancement of DENV infection, signifying that the presence of DENV E reactive IgA (especially fusion loop reactive IgA) has the potential to significantly modulate DENV infection and associated immunopathogenesis.

## Discussion

In this study we demonstrate that DENV-reactive IgA monoclonal antibodies can bind and neutralize DENV but are incapable of facilitating ADE of DENV infection *in vitro*. Furthermore, the presence of DENV-reactive IgA can significant blunt the DENV-infection enhancing activity of both DENV-reactive monoclonal IgG and polyclonal DENV-immune serum in a completive fashion. These results suggest an unappreciated role for DENV-reactive IgA during the humoral response to DENV infection and raise the potential that IgA could act as either a natural or therapeutic regulator of DENV dissemination and infection-attendant inflammation.

Although we have shown that DENV-reactive IgA is capable of disruption IgG mediated ADE, IgA may not be unique in this respect. Indeed, the depletion of IgM from flavivirus-immune serum has been shown to increase antibody-dependent enhancement of Zika virus infection of K562 cells, presumably by removing IgM as an antagonist of IgG-mediated infection enchantment [36]. Accordingly, parallel lines of evidence suggest that the ability of DENV-reactive IgG to interact with FcγRs is linked to the infection-enhancing potential of DENV-immune serum and – by extension – the clinical severity of DENV infection. Polymorphisms in FcγRIIa – a component of the low-affinity IgG receptor complex – have been associated with a decreased likelihood of becoming either symptomatically infected or progressing to severe dengue after DENV exposure [37-39]. While the mechanism behind the phenomenon is hasn’t been definitively established, it has been shown that at least some of these polymorphisms decrease the affinity of FcγRIIa for IgG [40]. Furthermore, post-translational modifications of IgG antibodies have been shown to significant impact their affinity for FcγR complexes and to correlate with dengue severity [41]. Most notably, the presence of high levels of afucosylated IgG– a post-translational modification which increases the affinity of the IgG Fc domain for FcγRIIIa [42] – either before or after DENV infection has been associated with increased dengue severity [43-45]. Finally, the presence of serum complement – such as C1q and C3 - can inhibit IgG mediated ADE both *in vitro* and *in vivo*, ostensibly by interfering with the ability of IgG Fc to interact with FcγR and/or forcing a complement-bound antibody into a configuration that is not amenable to fusion and viral entry [46-48].

While not found in nature, abolishing the ability of IgG antibodies to interact with FcγR through genetic engineering the Fc portion of IgG (LALA mutation) also ablates many of the infection-enhancing properties of DENV-reactive IgG both *in vitro* [35] and *in vivo* [49, 50]. Collectively, these this evidence underlines the importance of IgG/FcγR interactions the process of ADE, and emphasize the potential diagnostic and therapeutic implications for factors that can disrupt this immunologic nexus.

The data presented herein demonstrate that DENV-reactive IgA is capable of antagonizing IgG mediated enhancement of DENV infection, yet it is still unclear what role this process plays *in vivo* during natural DENV infection. Several previous studies have noted that high levels of serum IgA are associated with more severe disease following secondary DENV infection [28-31]. However, we and others have observed a higher frequency of IgA expressing plasmablasts following uncomplicated primary DENV infection than following severe secondary DENV infections [32, 33]. A key takeaway from the analysis performed in this study is that the absolute concentration of a given DENV-reactive antibody isotype may be a incomplete indicator of the infection-enhancing potential of a serum sample. Severe dengue is accompanied by robust production of antibodies of all isotypes, so without additional context the absolute level of any given single antibody isotype may provide an incomplete or misleading impression of immunologic features associated with disease severity.

## Materials and Methods

### Viruses

DENV-3 (strain CH53489) propagated in Vero cells were utilized for ELISA, FlowNT50, and ADE assays. Virus for ELISA was purified by ultracentrifugation through a 30% sucrose solution and the virus pellet was resuspended in PBS.

### Cell lines

Human K562 cells were maintained in IMDM supplemented with 10% FBS, penicillin, and streptomycin. U937-DC-SIGN cells were maintained in RPMI supplemented with 10% FBS, L-glutamine, penicillin, and streptomycin.

### Monoclonal antibodies and serum

The variable regions from the heavy and light chains were codon optimized, synthesized *in vitro* and subcloned into a pcDNA3.4 vector containing the human IgG1 or IgA1 Fc region by a commercial partner (Genscript). Transfection grade plasmids were purified by maxiprep and transfected into a 293-6E expression system. Cells were grown in serum-free FreeStyle 293 Expression Medium (Thermo Fisher), and the cell supernatants collected on day 6 for antibody purification. Following centrifugation and filtration, the cell culture supernatant was loaded onto an affinity purification column, washed, eluted, and buffer exchanged to the final formulation buffer (PBS). Antibody lot purity was assessed by SDS-PAGE, and the final concentration determined by 280 nm absorption. The clonotype information for all monoclonal antibodies generated as part of this study is listed in **Table 1**. Dengue IgG antibody positive plasma was purchased from SeraCare. Donor ID and batch numbers are shown in **Supplemental Table 1**.

### DENV-capture ELISA

Monoclonal antibody and plasma DENV-reactivity was assessed using a 4G2 DENV capture ELISA protocol. In short, 96 well NUNC MaxSorb flat-bottom plates were coated with 2 μg/ml flavivirus group-reactive mouse monoclonal antibody 4G2 (Envigo Bioproducts, Inc.) diluted in borate saline buffer. Plates were washed and blocked with 0.25% BSA + 1% Normal Goat Serum in PBS after overnight incubation. DENV-3 (strain CH53489) diluted in blocking buffer was captured for 2 hr, followed by extensive washing with PBS + 0.1% Tween 20. Serially diluted monoclonal antibody samples were incubated for 1 hr at RT on the captured virus, and DENV-specific antibody binding quantified using anti-human IgG HRP (Sigma-Aldrich, SAB3701362). Secondary antibody binding was quantified using the TMB Microwell Peroxidase Substrate System (KPL, cat. #50-76-00) and Synergy HT plate reader (BioTek, Winooski, VT). Antibody data were analyzed by nonlinear regression (One site total binding) to determine EC50 titers in GraphPad Prism 8 (GraphPad Software, La Jolla, CA).

### Neutralization Assay

Neutralizing titers of monoclonal antibodies and heat-inactivated plasma were assessed using a flow cytometry-based neutralization assay in U937 cells expressing DC-SIGN as previously described [51, 52]. Four-fold dilutions of antibody or sera were mixed with an equal volume of virus diluted to a concentration to achieve 10%–15% infection of U937-DC-SIGN cells in the absence of antibody. The antibody/virus mixture was incubated for 1 h at 37 °C, after which an equal volume of medium (RPMI-1640 supplemented with 10% FBS, 1% penicillin/streptomycin, 1% l-glutamine (200 mM) containing 5 × 10^4^ U937-DC-SIGN cells was added to each well and incubated 18–20 hr overnight in a 37 °C, 5% CO2, humidified incubator. Following overnight incubation, the cells were fixed with IC Fixation Buffer (Invitrogen, 00-82222-49), permeabilized using IC Permeabilization Buffer (Invitrogen, 00-8333-56) and immunostained with flavivirus group-reactive mouse monoclonal antibody 4G2 (Envigo Bioproducts, Inc.), and secondary polyclonal goat anti-mouse IgG PE-conjugated antibody (#550589, BD Biosciences). The percentage of infected cells were quantified on a BD Accuri C6 Plus flow cytometer (BD Biosciences). Data were analyzed by nonlinear regression to determine 50% neutralization titers in GraphPad Prism 8 (GraphPad Software, La Jolla, CA).

### ADE Assay

*In vitro* antibody-dependent enhancement (ADE) of DENV-3 infection was quantified as previously described [32, 53]. Four-fold serial dilutions of antibody or heat-inactivated sera were incubated with virus (in sufficient amounts to infect 10%–15% of U937-DC-SIGN cells) at a 1:1 ratio for 1 h at 37 °C. This mixture was then added to a 96-well plate containing 5 × 10^4^ K562 cells per well in duplicate. Cells were cultured for 18–20 hr overnight in a 37 °C, 5% CO2, humidified incubator. Processing and quantification continued as outlined in the FlowNT50 methods.

### Statistical Analysis

All statistical analysis was performed using GraphPad Prism 8 Software (GraphPad Software, La Jolla, CA). A P-value□<□0.05 was considered significant.

## Disclaimer

The opinions or assertions contained herein are the private views of the authors and are not to be construed as reflecting the official views of the US Army or the US Department of Defense. Material has been reviewed by the Walter Reed Army Institute of Research. There is no objection to its presentation and/or publication.

