## Supplemental figures for "Monomeric IgA antagonizes IgG-mediated enhancement of DENV infection"

### Supplemental Figure 1: Neutralization assay gating strategy

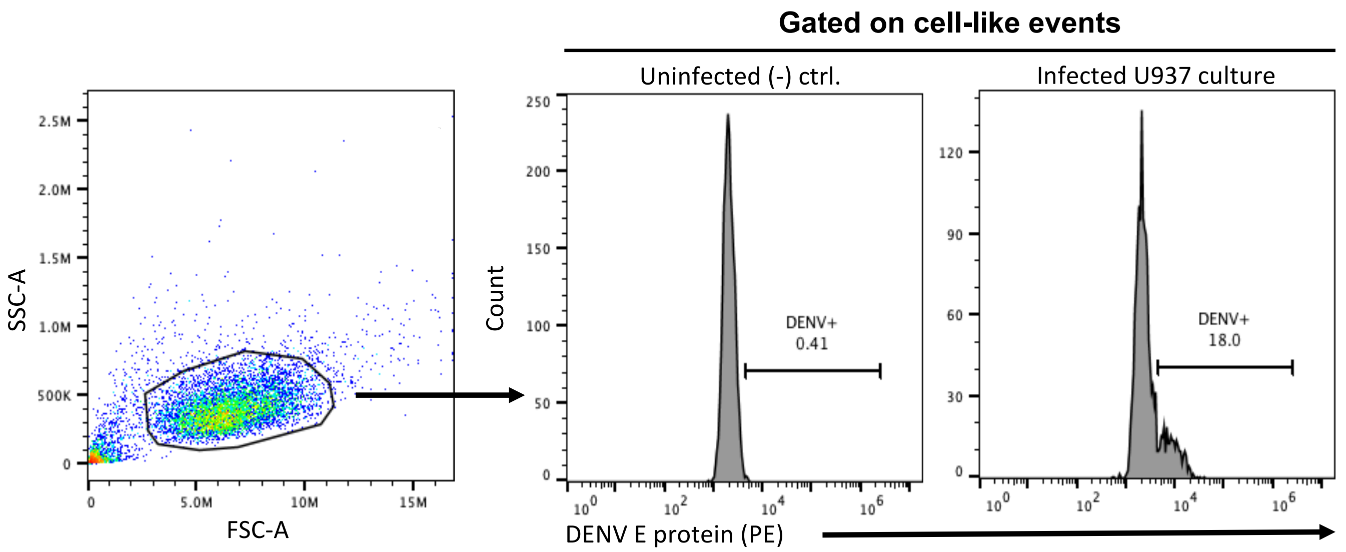


**Supplemental Figure 1.** Gating scheme and **r**epresentative plots from FlowNT assays

#### Supplemental Figure 2: ADE assay gating strategy

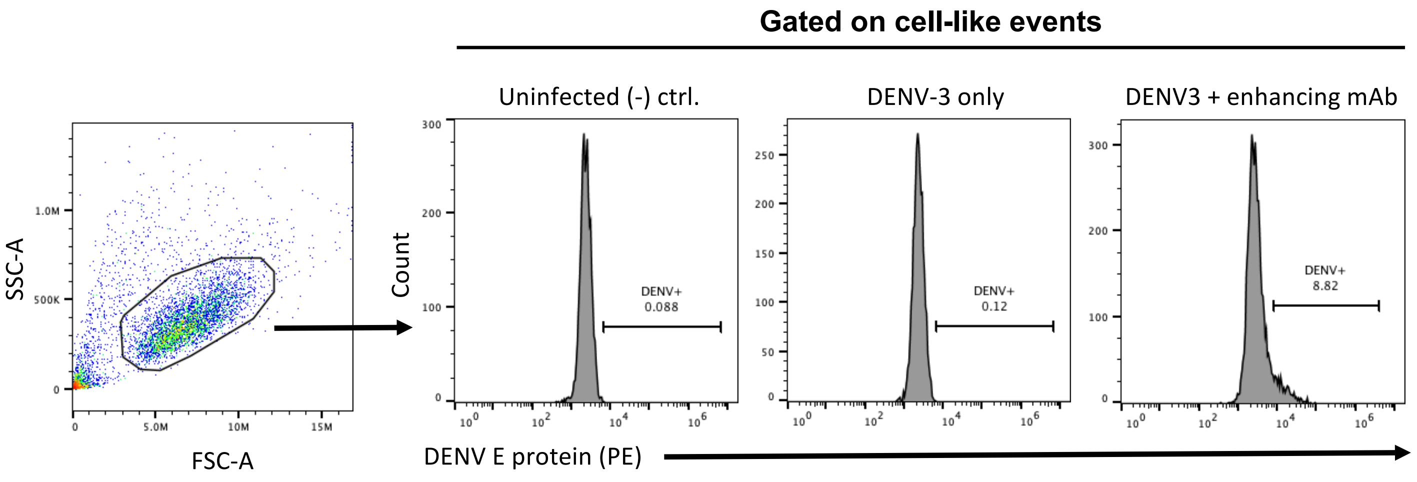


**Supplemental Figure 2.** Gating scheme and **r**epresentative plots from ADE assays

### Supplemental Figure 3: ELISA for IgM/IgG/IgA in DENV-immune plasma samples

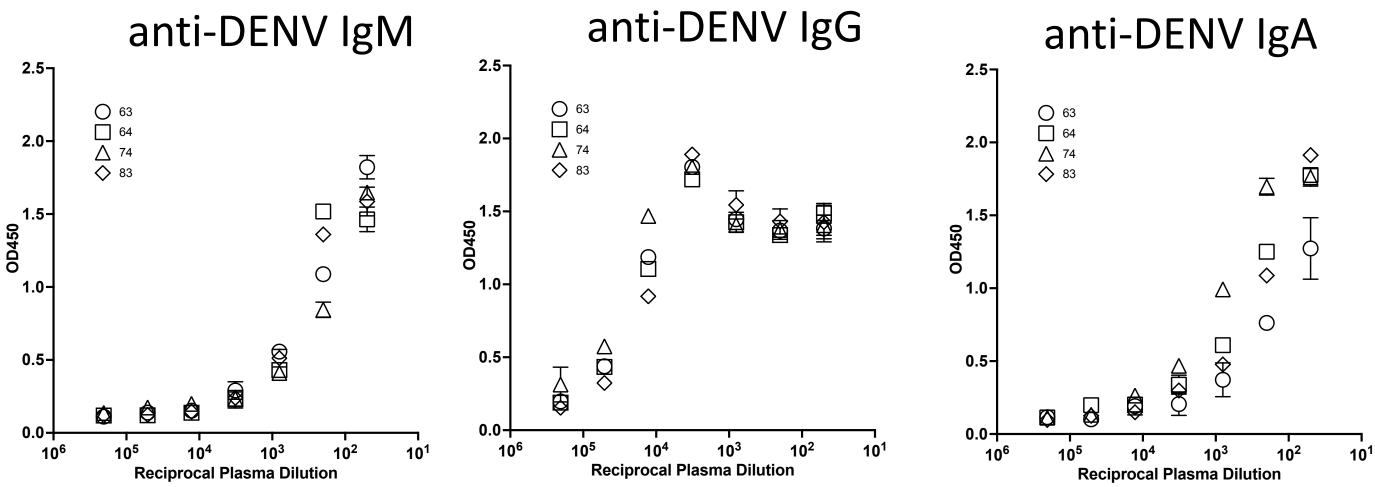


**Supplemental Figure 3.** DENV-3 IgM, IgG, and IgA titers in DENV-immune plasma samples. All analyzed samples exhibited significant DENV-3 IgG titers and modest IgM/IgA titers. Error bars +/- SEM

#### Supplemental Figure 4: DENV neutralization and ADE capacity of DENV-immune plasma samples

**A)**

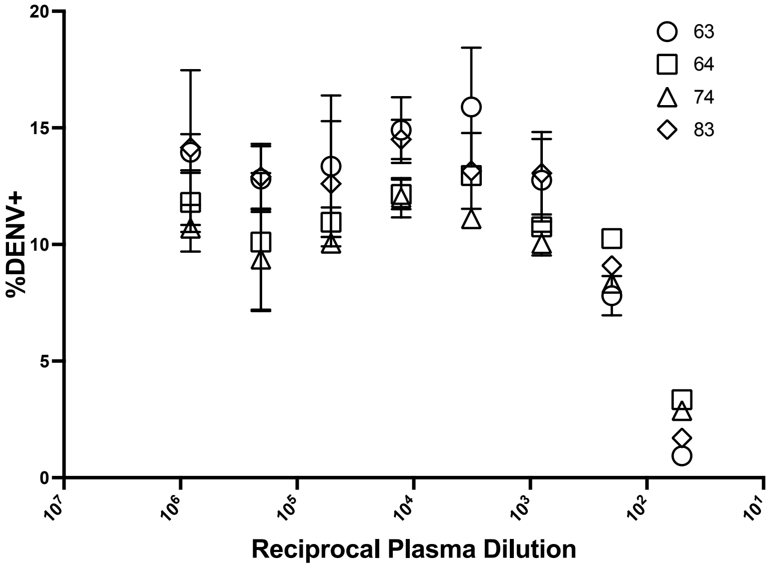


**B)**

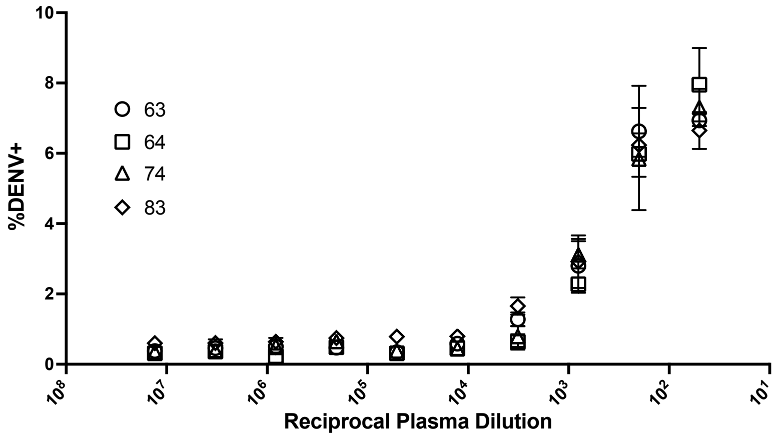


**Supplemental Figure 4. A)** DENV-3 neutralization activity of DENV-immune plasma as assessed by FlowNT. **B)** DENV-3 ADE activity of DENV-immune plasma as assessed by K562 infection. Error bars +/- SEM
