## Supplemental table for "Monomeric IgA antagonizes IgG-mediated enhancement of DENV infection"

**Supplemental Table 1. Dengue immune plasma used in this study**

| **Supplier** | **Product #** | **Donor ID** | **Batch number** |
| --- | --- | --- | --- |
| SeraCare | 0325-0014 | BD250524 | 10127363 |
| SeraCare | 0325-0014 | BD250525 | 10127364 |
| SeraCare | 0325-0014 | BD250535 | 10127374 |
| SeraCare | 0325-0014 | BD250543 | 10127383 |
